# Combating citrus greening disease by simultaneously targeting the pathogen and callose-mediated phloem occlusion

**DOI:** 10.64898/2026.01.03.697484

**Authors:** Maryam Khalilzadeh, Chun-Yi Lin, John M. Chater, Amit Levy, Christopher Vincent

## Abstract

Huanglongbing (HLB), caused by *Candidatus* Liberibacter asiaticus (*C*Las), disrupts sugar transport in citrus due to bacterial proliferation and excessive callose deposition that blocks sieve elements. Unblocking phloem may reduce symptoms, but it is unclear whether targeting *C*Las or suppressing callose deposition is more critical, as both contribute to phloem dysfunction. This study evaluated whether an integrated strategy targeting both *C*Las and callose-mediated phloem blockage could more effectively restore phloem transport, carbohydrate allocation, and productivity in HLB-affected citrus. Two complementary field experiments assessed the effects of an antibiotic, a callose inhibitor, and their combination on phloem recovery and source-sink dynamics. The first experiment focused on treatment-induced changes in carbon transport between source and sink shortly after delivery via stem infiltration, capturing immediate physiological responses. The second experiment assessed cumulative carbon allocation to determine sustained effects on assimilate transport and fruit development. Callose inhibitor treatment increased sugar transport but this immediate response did not yield lasting benefits. Antibiotic treatment enhanced carbon import into developing fruits, but these effects were not consistently supported by other physiological indicators. In contrast, the combined treatment reduced callose deposition, improved phloem conductivity and carbon allocation during high sink demand, decreased fruit abscission, and increased the total carbon accumulated in the fruits after physiological fruit drop. Collectively, these results demonstrate that impaired source-sink balance in HLB-affected citrus results from the combination of both pathogen load and host callose accumulation, supporting an integrated strategy that combines pathogen suppression with modulation of host phloem function to improve orchard productivity.

## Introduction

Huanglongbing (HLB; citrus greening) is currently the most destructive disease affecting global citrus production. It is associated with infection by the phloem-restricted α-proteobacterium *Candidatus* Liberibacter asiaticus (*C*Las), transmitted by the Asian citrus psyllid *Diaphorina citri* (Hemiptera: Liviidae) (Bendix and Lewis 2018; Bové 2006). The disease has devastated major citrus-producing regions; in Florida alone, HLB resulted in more than 7.8 billion in cumulative economic losses between 2007 and 2020 (Graham et al. 2020).

A growing body of evidence shows that the physiological basis of HLB symptoms is closely linked to phloem impairment, particularly the occlusion of sieve elements by pathogen-associated callose deposition (Etxeberria et al. 2009; Koh et al. 2012). During *C*Las colonization, infected sieve elements show pronounced callose accumulation at sieve plates, collapse of the phloem architecture, and deformation of vascular bundles (Deng et al. 2019; Etxeberria et al. 2009; Etxeberria and Narciso 2015; Kim et al. 2009). Callose deposition is a canonical early defense response in plants, triggered by pathogen perception and aimed at restricting pathogen spread by forming papillae or plugging sieve pores (Chen and Kim 2009; Kauss 1996). In the case of HLB, however, this defensive callose response becomes overactivated, and the resulting excessive deposition contributes to phloem collapse, thereby intensifying disease severity (Folimonova and Achor 2010; Granato et al. 2019).

Beyond callose-mediated occlusion, phloem dysfunction can also stem from the proliferation of *C*Las within phloem tissues. Although Kim et al. (2009) reported no visible *C*Las aggregates in leaf midrib phloem, substantial bacterial accumulation has been consistently observed in sink tissues. For example, Achor et al. (2020) and Hilf et al. (2013) documented dense *C*Las populations within the sieve elements of seed coat vasculature, and Fang et al. (2021) similarly showed that *C*Las reaches exceptionally high densities in fruit phloem, coinciding with severe phloem collapse.

*C*Las population and callose accumulation in the phloem cells resulted in decreased sugar transport. Leaves with higher *C*Las titers had greater callose accumulation and a marked decline in sucrose transport, establishing a direct link between bacterial density and phloem blockage (Welker et al. 2022). Impaired export of photoassimilates leads to carbohydrate accumulation in source leaves and pronounced starch buildup. This accumulation triggers negative feedback on photosynthetic capacity, evidenced by reduced CO₂ assimilation and down-regulation of photosynthesis-related proteins, ultimately producing widespread metabolic imbalance and foliar symptom development (Cai et al. 2022; Keeley et al. 2022; Koh et al. 2012; Ma et al. 2022; Nwugo et al. 2013). Consequently, HLB-affected trees show severe disruptions in source-sink dynamics: developing fruit receive insufficient sugars, leading to premature fruit drop, reduced fruit size and misshapen fruit, diminished juice quality, and overall declines in yield (Bassanezi et al. 2011; Rosales and Burns. 2011; Tang et al. 2020).

Collectively, these findings indicate that phloem functional decline in HLB-affected citrus, caused by both excessive pathogen-induced callose deposition and bacterial aggregation within the sieve elements, forms the central physiological basis for impaired carbohydrate transport and symptom development. Despite this dual contribution, current management strategies focus predominantly on antimicrobial approaches, particularly oxytetracycline trunk injection, which can reduce *C*Las titers and lessen visible symptoms (Archer et al. 2023; Vincent et al. 2022). However, yield responses to oxytetracycline remain inconsistent, varying with cultivar, injection method, tree age, timing, and rootstock-scion combinations (Albrecht et al. 2025; Archer et al. 2022; Worbington et al. 2025). Hu and Wang (2016) further showed that substantial reductions in *C*Las titers do not always translate into measurable yield gains, underscoring that bacterial suppression alone is insufficient to restore phloem function. These inconsistent outcomes indicate that reducing the population of the pathogen may not fully resolve the underlying physiological impairment.

In parallel, efforts targeting callose accumulation have shown potential for mitigating HLB-induced phloem dysfunction. Field and molecular studies using chemical callose inhibitors or callose-related gene knockdown have demonstrated reductions in callose deposition and starch accumulation, along with modest improvements in canopy condition and fruit performance. For example, Li et al. (2016) reported that treatment with callose inhibitor, including 2-deoxy-D-glucose (2-DDG), slowed disease progression and partially improved yield and fruit quality in sweet orange (*Citrus* x *sinensis*) and mandarin (*Citrus reticulata*). At the molecular level, Yao et al. (2023) showed that knockdown of the callose synthase gene *CsCalS11* decreased callose deposition and starch accumulation in leaves of *C*Las-infected trees, indicating that targeting callose can alleviate phloem blockage. While these studies highlight the promise of callose inhibition, consistent large-scale evidence linking this approach to reliable yield gains is lacking. Moreover, strategies targeting callose alone do not address the bacterial populations responsible for phloem dysfunction.

Given the interconnected roles of pathogen proliferation and host-driven callose deposition in vascular disruption, approaches focusing on only one component may not fully mitigate HLB symptoms. Therefore, there is a need to evaluate integrated strategies that simultaneously reduce *C*Las populations and prevent or reverse callose-mediated phloem plugging.

We hypothesized that simultaneously targeting *C*Las and callose-mediated phloem occlusion would more effectively restore phloem transport and carbon allocation in HLB-affected citrus trees than single-target interventions. To test this hypothesis, we evaluated the individual and combined effects of an antibiotic and a callose inhibitor on *C*Las titers, callose accumulation in leaf phloem tissues, and carbohydrate translocation from source leaves to developing fruit. We further assessed sink activity and cumulative fruit carbon gain at the whole-tree scale to determine whether improvements in phloem function translate into restored source-sink relationships and enhanced fruit development. Collectively, this study demonstrates that dual-targeting strategy addressing both pathogen burden and host-induced phloem blockage provides greater physiological recovery than previously reported single-target treatments.

## Methods

### Materials and Methods

Two independent field experiments were carried out to assess and compare the impact of a callose inhibitor, an antibiotic, and their combination on HLB-affected citrus trees, with a particular focus on the phloem recovery, carbon movement and source-sink dynamic. The first field experiment was a two-week study designed to examine how these treatments affect the dynamic of carbon transport between source (leaf) and sink (fruit) shortly after treatment infiltration, which shows immediate physiological effects. The second field experiment, conducted over two consecutive years, evaluated cumulative carbon allocation to assess longer-term effects of treatment on assimilate transport and fruit development.

The first experiment was conducted on Five-year-old ‘Duncan’ grapefruit (*Citrus × paradisi*) trees in Haines City, FL, USA (28°08’35.8"N 81°42’11.4"W), while the second experiment used eight-year-old ‘OLL-8’ sweet orange (*Citrus × sinensis*) trees grown at the experimental citrus grove of the University of Florida, Citrus Research and Education Center in Lake Alfred, FL (28°06’54.3"N 81°42’45.0"W).

For this study, we employed oxytetracycline (OTC) as the antibiotic and two well-established callose inhibitors. The first was 2-deoxy-D-glucose (DDG), a common analog that competes with glucose, the natural substrate of callose synthase, and has been demonstrated to suppress callose accumulation triggered by both abiotic and biotic stresses (Bayles et al. 1990; Li et al. 2012; Radford et al. 1998). The second inhibitor used was 3-aminobenzamide (3AB), which has been shown to selectively block callose deposition induced by bacterial flagellin and powdery mildew fungi, but not by abiotic stresses, suggesting that it acts specifically against pathogen-stimulated callose formation (Keppler et al. 2018).

### 1 Short-term Field Experiment

#### Plant material and design

A total of 24 ‘Duncan’ grapefruit trees naturally infected with *C*Las were selected and randomly assigned to four treatments (six trees per treatment) following a completely randomized design. The four treatments were (1) OTC (ReMedium TI^®^ ; TJ BioTech LLC, Buffalo, SD, USA; 95.0% oxytetracycline hydrochloride) 1 mg ml^-1^, (2) callose inhibitor (DDG) 0.007 mg ml^-1^, (3) combined solution containing both OTC and DDG, and (4) no treatment.

#### Infiltration procedure

Two stems per tree, each bearing one fruit and several leaves, were selected. The leaves were defoliated to have only three leaves per stem to have consistency among treatments and replicates. The stems were girdled by cutting a ring of bark near the base and wrapping the girdle with parafilm. A latex glove finger (cut open on both sides) was placed over the stem to create a small infiltration chamber. Small punctures were made around the stem using a dermal roller (0.5-1.5 mm spikes). One milliliter of infiltration solution was added to each infiltration chamber, which was then sealed with parafilm and secured with a zip tie to prevent leakage (Supplementary Figure 1). Infiltration process was carried out in the middle of April 2024.

#### Sucrose-¹⁴C labeling

Two weeks after infiltration, stems bearing a single fruit were excised from the girdled area, transported to the laboratory, and prepared for sucrose-¹⁴C labeling. The ends of the stems were placed in Eppendorf tubes filled with distilled water, and the fruit was wrapped in parafilm to minimize transpiration. The nearest leaf to the fruit was abraded with sandpaper, fed with sucrose [¹⁴C(U)] (5 μCi) (Revvity, Waltham, MA, USA), and covered with aluminum foil for optimal absorption, with a retention time of 24 hours. Fruits, stems, and leaves were collected separately, lyophilized, and analyzed for ¹⁴C distribution using liquid scintillation counting as described in Li et al. (2024) (Supplementary Figure 1).

### 2. Two-year Field Experiment

#### Plant material and experimental design

The experimental trees were eight years old ‘OLL-8’ sweet orange (*C. sinensis*) grafted onto UFR-1, a tetraploid hybrid rootstock with a complex background of *C. reticulata*, *C. grandis*, and *Poncirus trifoliata* selected by UF/IFAS for improved tolerance to HLB (Grosser 2015). All trees were already naturally infected by *C*Las prior to our experiments. Treatments included a callose inhibitor (3AB), OTC and their combination, imposed in a randomized complete block design (n = 12 biological replicates per treatment). The trees were exposed to similar natural environment (air temperature, humidity, etc.) and were drip-irrigated daily and fertilized periodically. In early May 2024, and middle of April 2025, the treatments trunk injected under sunny conditions with an average midday temperature of approximately 28 °C. We applied OTC at its recommended concentration of 5,500 ppm and delivered the callose inhibitor at a concentration of 1 mM. Each tree received 50 mL of the prepared solution. Injections were performed on the north side of the trunk using a large FlexInject™ injector fitted with a 17/64-inch sharp brad-point drill bit (Supplementary Figure 2). Control trees received an equivalent injection of 50 mL of double-distilled water, applied to the same number of trees as the treatment group. Treatments were applied during mid- to late morning (10:00 a.m.–12:00 p.m.), with formulations freshly prepared on the morning of application.

#### Growth measurements

Canopy volume and density were recorded quarterly for 18 months following the first year of trunk injection (April 2024-October 2025). Canopy volume was estimated using a modified prolate spheroid equation. Tree height (H) and canopy diameters in the east-west (D1) and north-south (D2) orientations were measured. Canopy volume (m³) was then calculated using the formula:

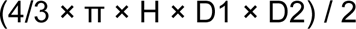

Canopy density was quantified using a Li-Cor LAI-2200 Plant Canopy Analyzer (Li-Cor, Lincoln, NE, USA) by taking measurements on the north and south sides of each tree, which were then averaged.

#### Pathogen assessment (*C*Las titer)

For each treatment, samples consisting of six young and six mature leaves were collected from 12 trees at the time of pre-injection and again at 2-, 4-, and 8-week post-injection. Leaves from three trees were combined, yielding four replicates per treatment. Midribs were excised, and total DNA was extracted using a modified CTAB method. Quantitative PCR (qPCR) was performed using primers targeting the rplJ/rplL ribosomal protein gene of *C*Las, following Wang et al. (2006). *C*Las titer was expressed as qPCR cycle threshold (Ct), with lower Ct indicating higher bacterial load. Trees were classified as *C*Las-positive if their Ct values were below 32. Ct values from 32 to 36 were considered weakly positive, while Ct values above 36 indicated a *C*Las-negative plant, consistent with thresholds used in previous *C*Las diagnostic studies (Alves et al. 2021; Coy et al. 2014; Hoffman et al. 2013; Whitaker et al. 2014; Wulff et al. 2020;).

#### Fruit measurements

On each tree, 30 fruits were tagged (Supplementary Figure 2), and their diameters were measured using a digital caliper before treatment application in mid-April (pre-injection) and then biweekly at five subsequent time points: end of April (2 weeks), mid-May (4 weeks), end of May (6 weeks), mid-June (8 weeks), and end of June (10 weeks post-injection). At each measurement date, three additional fruits per tree were collected to determine fruit diameter and dry weight. For dry weight measurements, fruits were cut and oven-dried for four days before being weighed. The resulting paired data of diameter and dry weight were used to develop a regression model to estimate the dry weight of the 30 tagged fruits, for which only diameter data were available. At each time point, the percentage of fruit drop relative to the 30 tagged fruits was also recorded.

Fruit collected in mid-May (4 weeks post-injection), and mid-June (8 weeks post-injection) were also used to measure dark respiration. Dark respiration was quantified using a custom-built chamber integrated with an LI-6800 infrared gas analyzer (LI-COR Biosciences Inc.). Instrument conditions were standardized during all measurements, including an airflow of 600 μmol s⁻¹, chamber pressure maintained at 0.1 kPa, and an internal fan speed set to 14,000 rpm, water vapor at 30 mmol, CO₂ concentration stabilized at 400 μmol mol⁻¹, and chamber temperature matched to ambient conditions. Before measurement, fruits were dark acclimated for 20 minutes inside black plastic bags to ensure the absence of photosynthetic activity. Three Fruits were placed in airtight chamber and measurements were conducted in darkness and therefore represented CO₂ efflux exclusively during the nocturnal period. The IRGA provided chamber-level CO₂ efflux rates in µmol CO₂ s⁻¹. First, CO₂ fluxes were converted from µmol to mol CO₂. CO₂ efflux was expressed on a per-gram basis by dividing chamber respiration by the total dry mass of the three fruits. This mass-specific rate was then multiplied by the dry mass of each fruit to assign an appropriate fraction of chamber respiration to each fruit. The total moles of CO₂ were expressed as mol glucose equivalents by dividing by six, corresponding to the complete oxidation of one mole of glucose releasing six moles of CO₂. Glucose equivalents were then calculated in grams by multiplying by the molar mass of glucose (180.156 g mol⁻¹). Finally, respiration rates (g glucose s⁻¹ fruit⁻¹) were scaled to the corresponding nighttime duration to estimate daily glucose loss per fruit.

#### Biochemical analysis

*Callose quantification*: Callose extraction, quantification and data analysis were performed following Bernardini et al. (2024). Six mature leaves per tree were collected, frozen, and processed for midrib tissue extraction. Callose was extracted from 20 mg of ground tissue and isolated using sequential ethanol washes, alkaline extraction, sonication, and heat treatment. The resulting supernatant was used for fluorescence-based quantification, with parallel measurements to correct for sample autofluorescence. Fluorescence was measured at 360 nm excitation and 460 nm emission using a fluorescence spectrophotometer (BioTek Fluorescence Microplate Readers FLx800), and callose content was normalized to fresh tissue mass (µg mg⁻¹ FW) following Bernardini et al. (2024).

*Starch quantification*: Leaf samples from the trees treated with the combination treatment were collected and flash frozen. Starch was extracted from 25 mg of dried ground tissue. Samples were boiled in Milli-Q water, centrifuged, and separated into supernatant and pellet fractions. Starch was quantified from the pellet after ethanol washes to remove impurities. Starch was enzymatically digested with amyloglucosidase and α-amylase, followed by colorimetric determination using the anthrone method. Absorbance was measured at 620 nm, and concentrations were calculated against standards.

#### Estimation of dry mass and dark respiration

Regression models were developed to predict dry mass and dark respiration of the tagged fruits attached to the trees, based on fruit diameter and subsampling for dry matter and respiration. Because dry matter and respiration required destructive sample, this approach was used to enable us to estimate the carbon accumulation and use of fruits that remained on the tree nondestructively. The first regression model was constructed to predict fruit dry weight (DW). The model incorporated the main effects of fruit diameter (FD), treatment, and time, as well as their interactions, to account for potential differences in the FD-DW relationship across treatments and developmental stages. To respect the experimental design, block was included as a random effect. The model was calibrated using the three destructively sampled fruits per tree collected at sampling timepoint, for which both FD and DW were measured. The final model was then applied to the 30 tagged fruits per tree, for which FD had been measured nondestructively, to estimate their DW.

The statistical model was specified as:

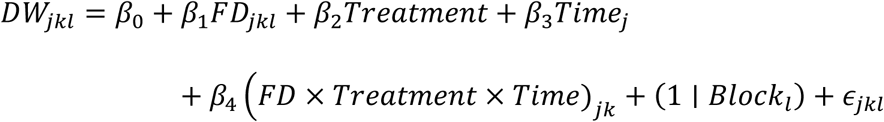

where 𝐷𝑊_jkl_is the dry weight of fruit 𝑘 from tree in block 𝑙 at time point 𝑗, *β₀* is intercept and 𝜖_jkl_ is the residual error. Model selection was performed using backward stepwise regression guided by Akaike’s Information Criterion (AIC), adjusted R² and p-values of interaction terms. At each step, nested models were compared using ANOVA, and the final model was chosen according to the principle of parsimony, retaining the simplest structure that adequately captured the variation in the data.

Fruit respiration was quantified as glucose loss per fruit per day, derived from dark respiration measurements. Paired data of respiration rate and fruit dry weight were used to construct an allometric regression model predicting respiration-derived glucose loss as a function of DW, while accounting for treatment and developmental stage. The model included log-transformed DW, treatment, and time (and their interactions) as fixed effects, with block treated as a random effect. Calibration was performed using destructively sampled fruits for which both respiration and DW were directly measured. The resulting model was then applied to the 30 tagged fruits whose DW values were estimated from diameter measurements to predict respiration rates per fruit.

#### Carbon-balance and total relative crop growth calculations

Carbon import was calculated using a simplified carbon-balance approach that integrates biomass gain with respiratory carbon losses. For each fruit the rate of dry-weight accumulation (𝑫𝑾𝒕𝟐-𝑫𝑾𝒕𝟏) was combined with the corresponding respiration rate measured over the same period. Because citrus fruit exhibit negligible net photosynthesis, the assimilation term in the full carbon-balance equation was assumed to be zero. The resulting expression, 𝑪import = 𝑫𝑾𝒕𝟐-𝑫𝑾𝒕𝟏 + 𝑹, provided an estimate of the net carbon imported into each fruit on a per-day basis.

Total relative crop growth was calculated as the cumulative carbon gain over time which expressed as grams of dry weight (gDW) and was determined by summing the carbon accumulated in 30 tagged fruits over a ten-week period following injection.

### 3. Statistical Analysis

All statistical analyses were performed in R (version 4.3.1). Data were checked for normality and homogeneity of variances prior to analysis, and log transformations were applied when necessary to meet assumptions. To assess treatment effects on sucrose-¹⁴C export, differences between each treatment and the non-treated control were evaluated using independent t-tests because of high within-group variability. Resulting p-values were adjusted using the Holm–Bonferroni correction method (Holm, 1979) to control the family-wise error rate. Sucrose-¹⁴C data were analyzed using beta regression (logit link) implemented in the R package betareg (Cribari-Neto & Zeileis, 2010), which is appropriate for modeling response variables expressed as continuous proportions ranging from 0 to 1. Total recovery fraction was included in the model to account for variation in labeling efficiency among samples.

Linear mixed-effects models were fitted using the lme4 package (lmer function; Bates et al., 2015), with treatment and time as fixed effects and block as random effect when applicable. Model significance was assessed using type III ANOVA (via the car package; Fox & Weisberg, 2018), and pairwise comparisons among treatments were conducted using estimated marginal means (emmeans) followed by post hoc tests with appropriate multiple-comparison adjustments. Linear regressions (lm) were used where linear mixed-effects models were not required. Data visualization and diagnostic plots were generated using ggplot2 (Wickham, 2011).

## Results

### 1. Short-term Field Experiment Carbon allocation following treatment

To assess the immediate physiological effects of callose inhibitor and antibiotic treatments on carbon transport in HLB-affected citrus trees, ¹⁴C-labeled sucrose was applied to source leaves two weeks after infiltration treatment. The recovered radioactivity was measured in leaf, stem, and fruit tissues, and the proportion of ¹⁴C distributed among these organs was compared among treatments. A beta regression model with a logit link was used to evaluate treatment effects on the proportion of ¹⁴C-labeled sucrose exported from source leaves and imported into fruits.

Infiltration with the callose inhibitor significantly increased both source export and sink import of ¹⁴C-labeled sucrose compared with the control, resulting in 2.20-fold (*P* = 0.01) and 2.81-fold (*P* < 0.001) increases, respectively. The combination of callose inhibitor and antibiotic produced positive but non-significant effects on sucrose transport, with 1.63-fold and 1.38-fold increases in export and import, respectively. In contrast, antibiotic treatment alone caused a modest, non-significant increase in export (1.44-fold) and a reduction in import (0.33-fold) (Fig 1).

**Fig. 1.**
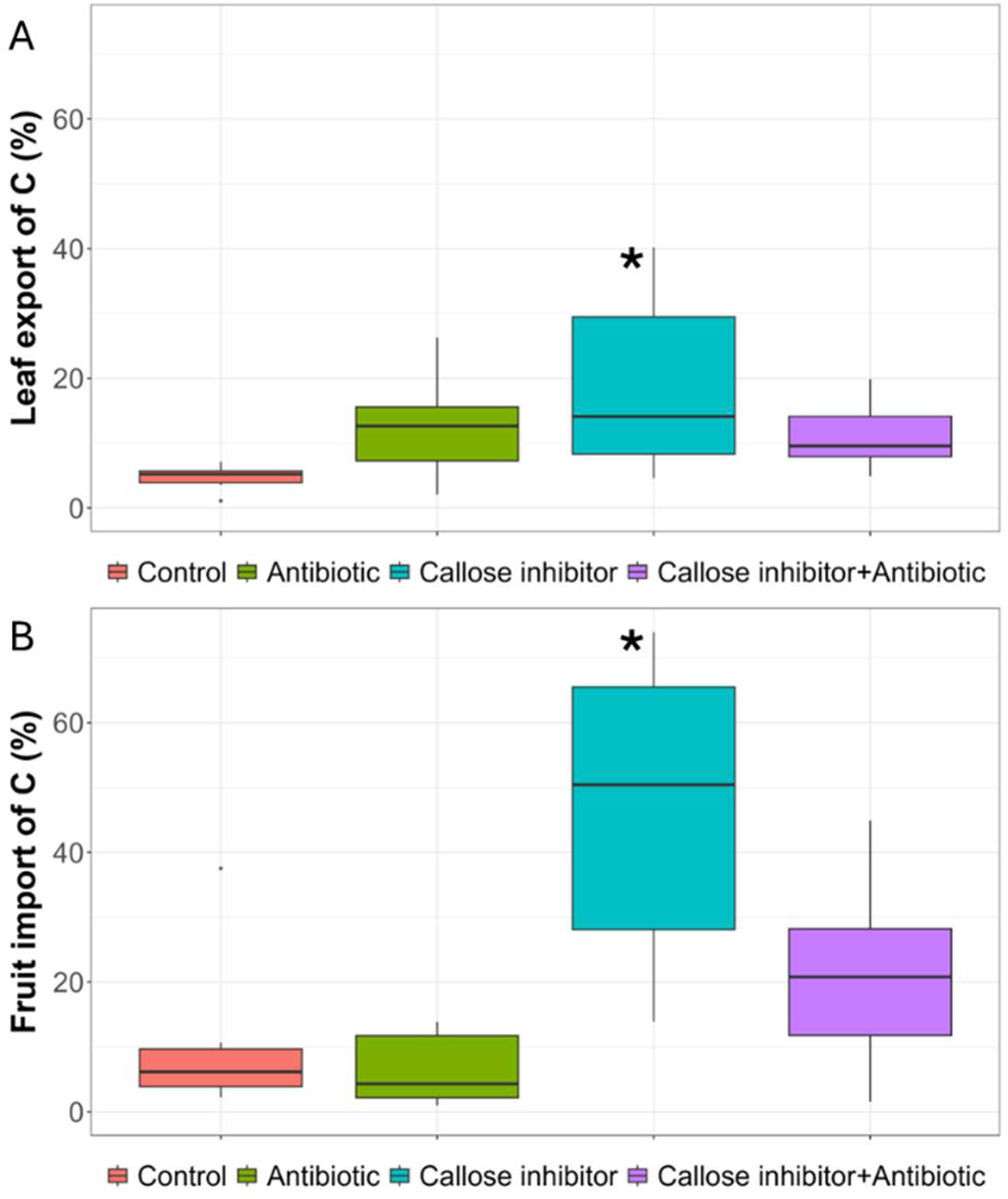
Treatment effects on dynamic of ¹⁴C-sucrose transport between source (leaf) and sink (fruit) two weeks post-infiltration. A) Percent total labeled ^14^C exported from leaf to stem and fruit and B) percent total labeled ^14^C imported to fruit of HLB-infected ‘Duncan’ grapefruit. Treatments consisted of 2-deoxy-D-glucose (DDG), oxytetracycline (OTC), a combined DDG + OTC solution, and an untreated control. ¹⁴C labeling was conducted on excised branches using ¹⁴C-sucrose under laboratory conditions. Asterisks denote significant differences from the control (*P* < 0.05).

### 2. Two-year Field Experiment

While reducing callose deposition may enhance sugar movement by reopening blocked phloem, thereby temporarily supporting tree function, over longer periods this reopening could also facilitate the movement of the HLB-associated bacterium within the tree. As a result, the long-term physiological response may reflect a tradeoff between these opposing effects. To understand how these factors play out over time, we next carried out a two-year field study on HLB-infected ‘OLL-8’ sweet orange trees that were trunk-injected with either antibiotic alone, callose inhibitor alone, or a combination of both.

#### Tree growth and canopy development

Relative growth rate (RGR) of canopy volume changed significantly over time, increasing from April 2024 to November 2024, dropping by April 2025, and rising again by October 2025 but it was unaffected by treatment (Fig 2A). The temporary decline following November 2024 coincided with a post hurricane event, which likely caused branch dieback, reducing canopy volume, suggesting that the observed changes in canopy volume were driven largely by environmental events, like hurricane damage, rather than treatment effects.

**Fig. 2.**
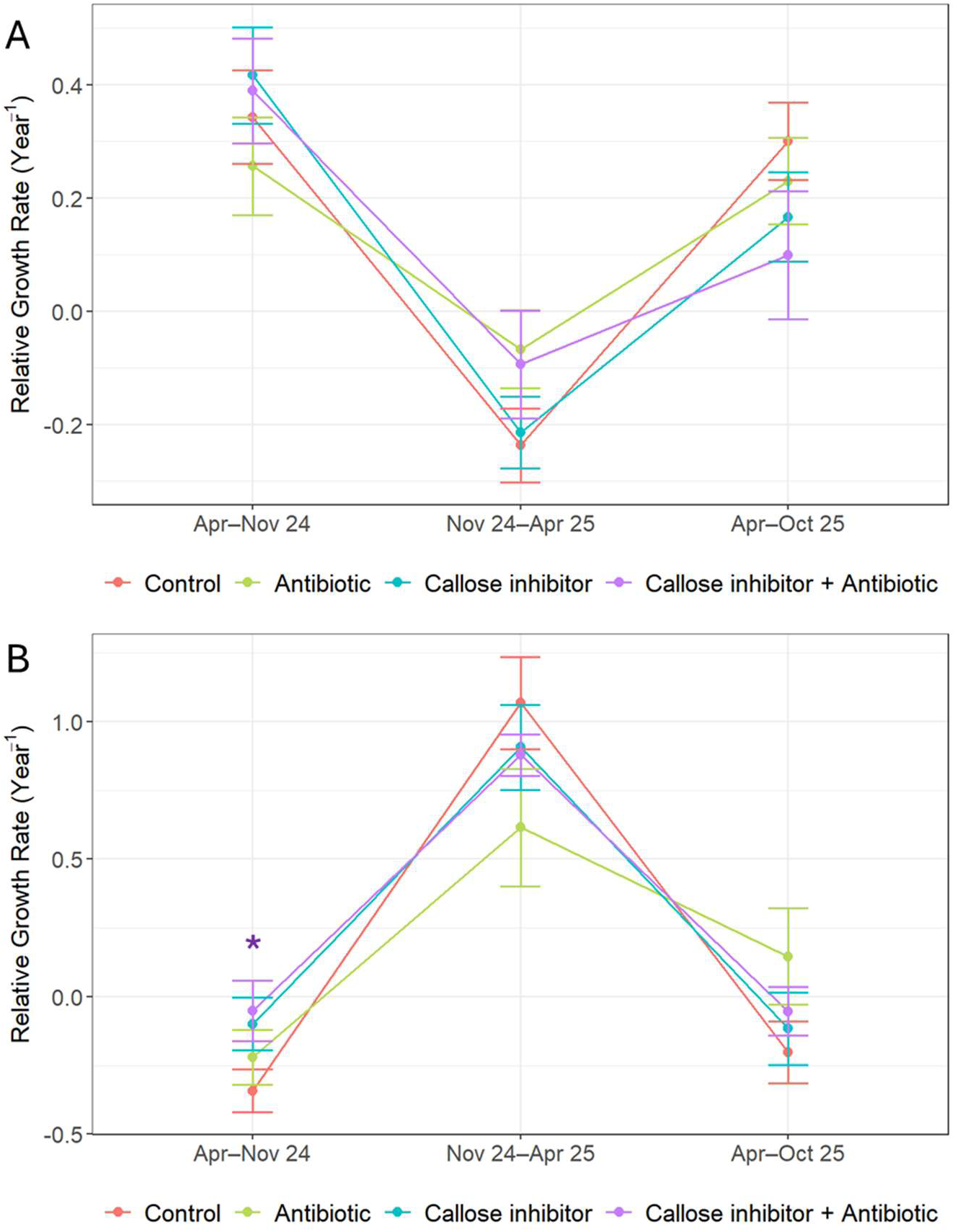
Effects of treatments on vegetative growth, including (A) Relative growth rate of canopy volume and (B) Relative growth rate of canopy density, in HLB-infected ‘OLL-8’ sweet orange (*C. sinensis*) grafted onto UFR-1 rootstock (*Citrus spp.*) over the 18-month from April 2024 to October 2025. Treatments included oxytetracycline, the callose inhibitor (3AB), their combined application, or untreated control. Asterisks, colored by treatment, denote significant differences from the control (*P* < 0.05).

The RGR of canopy density varied significantly over time, with the interval × treatment interaction approaching significance (*P* = 0.07). From April to November 2024, the callose inhibitor and combination treatments exhibited the smallest reductions in canopy density, suggesting greater canopy resilience during this period. This interval included the October hurricane (Hurricane Milton, October 9-10, 2024), which likely contributed to variations in canopy density. During this period, the combination treatment resulted in a significantly higher RGR of canopy density compared with the control (*P =* 0.039), while the callose inhibitor treatment showed a marginally higher RGR (*P =* 0.082). In contrast, the OTC treatment did not show a consistent effect on canopy density relative to the control, with no difference in the first interval (*P =* 0.378), a marginal decrease in the second interval (*P =* 0.053), and a marginal increase in the third interval (*P =* 0.066). Across the full 18-month study period, the combination treatment maintained a higher overall RGR of canopy density than the control (*P =* 0.069) (Fig 2B), indicating a sustained treatment effect under prolonged disease pressure.

#### Infection-level assessment

Overall, the Ct values for most treatments stayed within the weak-positive range, with a few exceptions. Weakly positive Ct values observed in trees showing clear HLB symptoms are likely a result of random whole canopy sampling, along with pronounced canopy-level variability and the uneven distribution of the pathogen within individual trees (Tatineni et al. 2008; Louzada et al. 2016). Before the injections, all groups showed Ct levels typical of weak-positive trees. At two and four weeks after treatment, the callose inhibitor group showed a clear drop in Ct values, placing those trees in the strong-positive category. By eight weeks, the trees that received the combination of callose inhibitor and antibiotic showed a noticeable rise in Ct values, moving them into the negative range (Fig 3). This improvement suggests that the combination treatment had a positive effect in reducing the tree infection level over time.

**Fig. 3.**
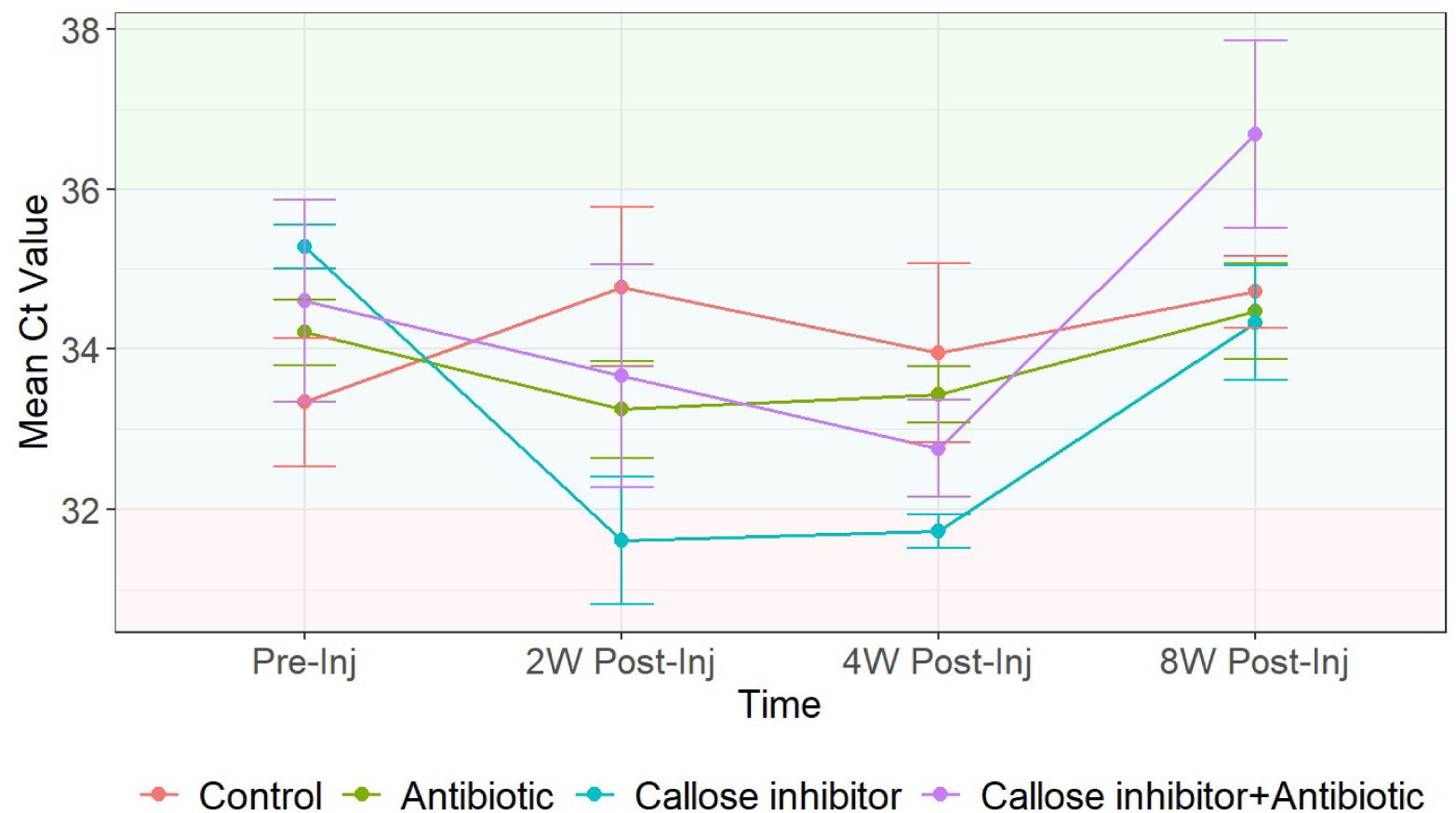
Cycle threshold (Ct) values obtained by qPCR for *Candidatus* Liberibacter asiaticus in HLB-infected ‘OLL-8’ sweet orange (*C. sinensis*) grafted onto UFR-1 rootstock (*Citrus spp.*), evaluated across four treatment groups (oxytetracycline, callose inhibitor (3AB), combined treatment, and control) and multiple sampling times including before injection (Pre-Inj) and at two, four, and eight weeks after injection (2W Post-Inj, 4W Post-Inj, and 8W Post-Inj), throughout the field trial. Color coding reflects Ct values categories: pink for *C*Las-positive (Ct < 32), blue for weakly positive (32 < Ct < 36), and green for *C*Las-negative (Ct > 36).

#### Fruit growth and retention dynamics

Treatment, interval, and their interaction all had significant effects on the relative fruit growth rate (Table 1). Trees receiving the combination treatment exhibited higher relative fruit growth rates during the initial two weeks following treatment injection and again during the six- to eight-week interval compared to the control. The relative fruit growth rate of callose inhibitor-treated trees was significantly lower during the four- to six-week interval compared to the control (Fig 4). Over the entire measurement period, absolute fruit growth was highest in trees receiving the combined treatment, while trees treated with the callose inhibitor alone showed the lowest absolute growth (Fig 4).

**Fig. 4.**
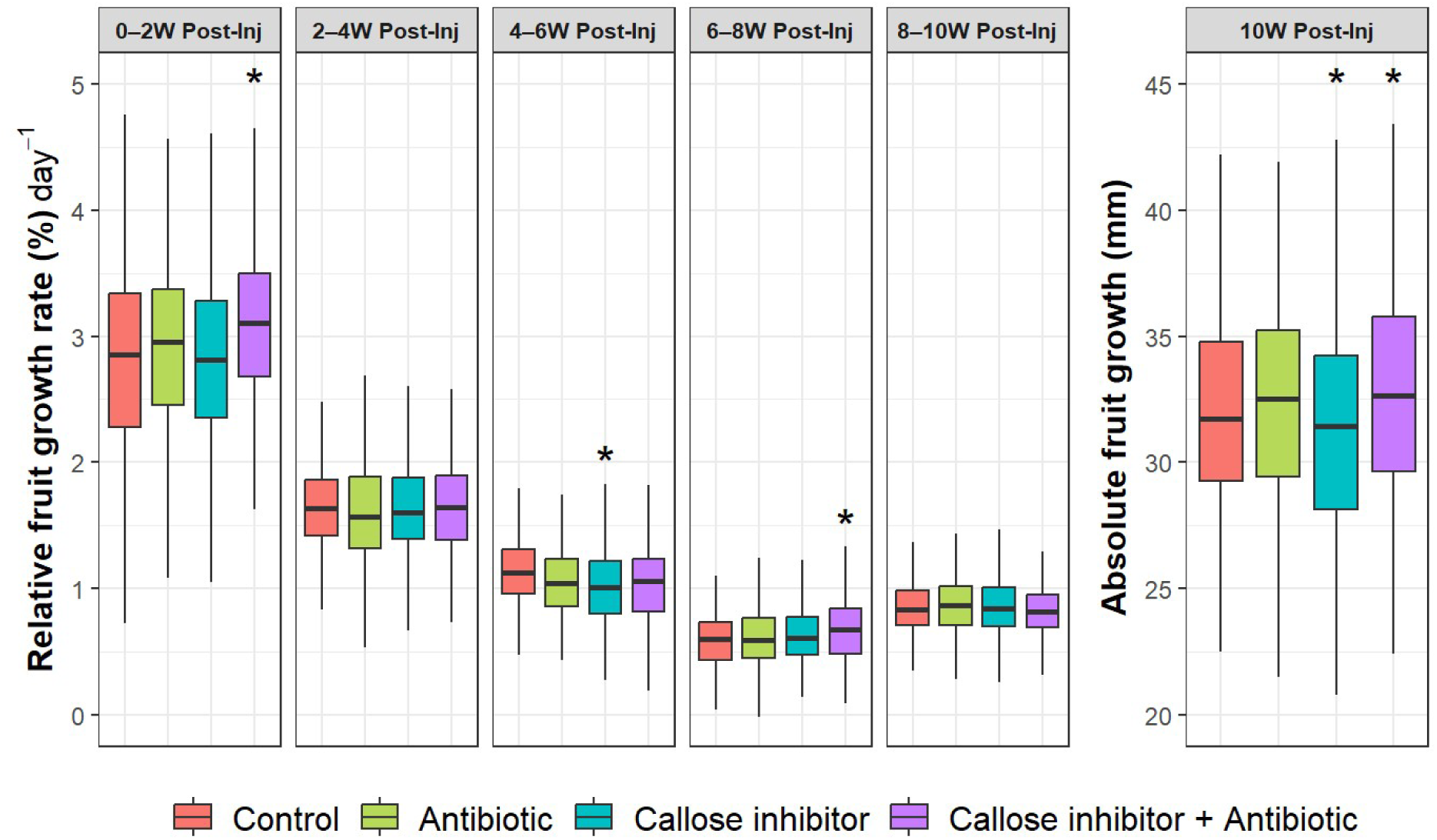
Absolute and Relative fruit growth rate of HLB-infected ‘OLL-8’ sweet orange (*C. sinensis*) grafted onto UFR-1 rootstock (*Citrus spp.*), treated with oxytetracycline, the callose inhibitor (3AB), their combination, or no treatment (control), measured across five post-injection intervals. Absolute growth represents the cumulative increase in fruit size over the entire measurement period, whereas relative growth reflects interval-specific changes. Asterisks indicate where treatments differ significantly from the control. Intervals: pre-injection to 2 weeks, 2-4 weeks, 4-6 weeks, 6-8 weeks, 8-10 weeks post-injection.

**Table 1.** Repeated-measures analysis of variance of relative fruit growth rate in HLB-infected ‘OLL-8’ sweet orange (*C. sinensis*) grafted onto UFR-1 rootstock (*Citrus spp.*), treated with oxytetracycline, the callose inhibitor (3AB), their combination, or left untreated (control) across five post-injection intervals.

| Effect | *df* | Sum Sq | *F*-value | *P*-value |
| --- | --- | --- | --- | --- |
| Treatment | 3 | 3.1 | 3.1582 | 0.0237 |
| Interval<sup>1</sup> | 4 | 3420.7 | 2594.1834 | < 0.0001 |
| Treatment × Interval | 12 | 18.3 | 4.6297 | < 0.0001 |

<sup>1</sup>Interval: pre-injection-2 weeks, 2-4 weeks, 4-6 weeks, 6-8 weeks, 8-10 weeks post-injection.

Both treatment and time independently had significant effects on physiological fruit drop (*P* < 0.0001), whereas their interaction was not significant (Fig 5).

**Fig. 5.**
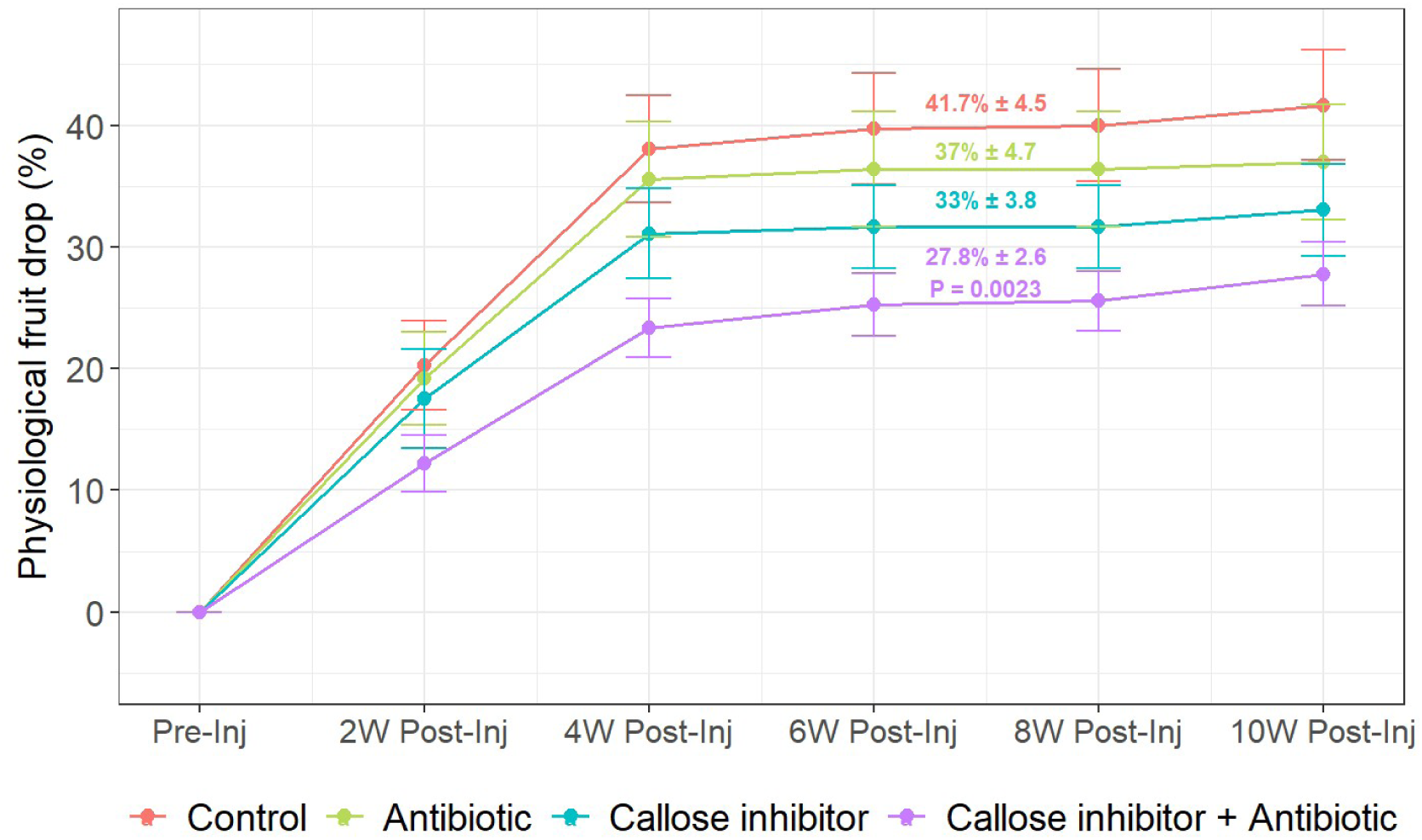
Proportion of physiological fruit drop in HLB-infected ‘OLL-8’ sweet orange (*C. sinensis*) grafted onto UFR-1 rootstock (*Citrus spp.*), treated at fruit set with oxytetracycline, callose inhibitor (3AB), their combination, or left untreated (control) across five post-injection intervals. Cumulative fruit drop was lowest in the combined treatment (27.8% ± 2.6) and was significantly lower than the control (41.7% ± 4.5; *P* = 0.0023), but not significantly different from the callose inhibitor (33% ± 3.8; *P* = 0.119) or antibiotic treatment (37% ± 4.7; *P* = 0.0855).

#### Fruit respiration, carbon allocation and total relative crop growth

Respiration was estimated for each of the 30 tagged fruits and used to evaluate how the treatments influenced fruit respiration. Carbon import was then calculated by combining each fruit’s dry-weight increase with its estimated respiratory carbon loss. The analysis of variance indicated that treatment, time, and their interaction had a significant effect on fruit respiration and carbon import into the fruits (*P* < 0.0001).

Four weeks after injection (mid-May), fruits from control trees showed the lowest glucose loss through respiration, while fruits under the combined treatment exhibited the highest loss, indicating slower metabolic carbon turnover in the control and higher turnover in the combined treatment, with intermediate responses in the other treatments. At eight weeks post-injection (mid-June), fruits from trees treated with antibiotic and the combination treatment showed greater respiration rates than those treated with callose inhibitor or control trees. Carbon import also differed among treatments and followed the same pattern observed for fruit respiration at both sampling times (Fig 6A).

**Fig. 6.**
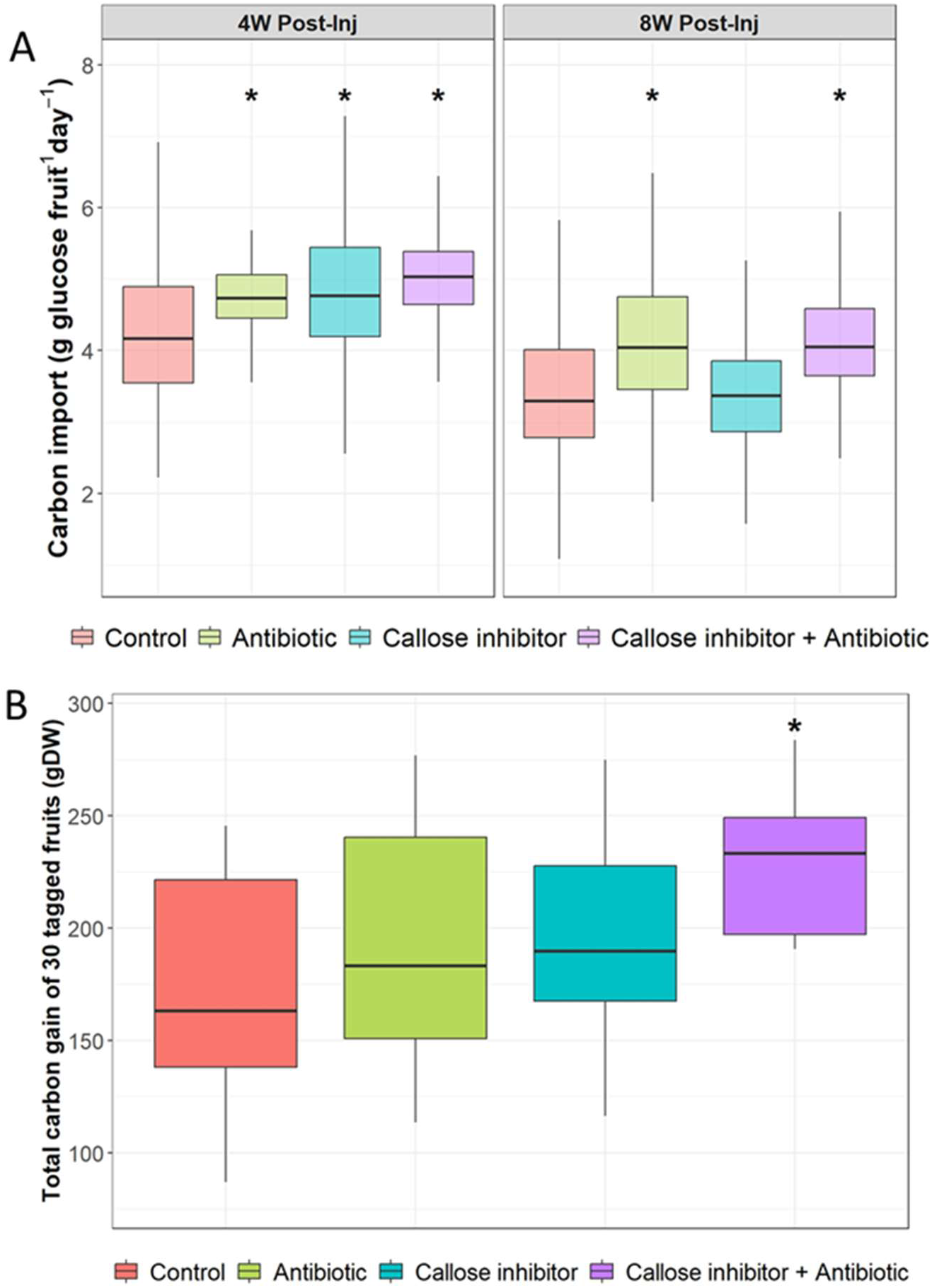
Effects of oxytetracycline, the callose inhibitor (3AB), their combined application, and untreated control on (A) carbon import into the fruit at four and eight weeks after trunk injection and (B) Total relative crop growth, expressed as the total carbon gain (gDW) from 30 tagged fruits summed over ten weeks after injection in HLB-infected ‘OLL-8’ sweet orange (*C. sinensis*) grafted onto UFR-1 rootstock (*Citrus spp.*). Treatments marked with an asterisk show a statistically significant increase relative to control.

Total relative crop growth was calculated as the total carbon gain (gDW) accumulated by 30 tagged fruits over ten weeks following injection and then compared among control trees and those receiving antibiotic, callose inhibitor, or combined treatments to assess treatment effects on fruit carbon accumulation, including accounting for fruit drop. The combination treatment resulted in the highest carbon gain, whereas the control was the lowest. The antibiotic and callose inhibitor treatments fell in between, showing slightly higher values than the control, but the differences were not statistically significant. This indicates that the combined application of the callose inhibitor and the antibiotic increases fruit carbon gain more effectively than either treatment alone when compared with the control (Fig 6B; *P* < 0.05).

#### Phloem recovery and source-sink balance

Callose levels in leaf midribs were significantly lower in trees treated with the callose inhibitor at 8 weeks after injection in 2024. Likewise, the combination treatments also resulted in reduced callose accumulation at the 8-week assessment in both 2024 and 2025 (Fig 7). Leaf starch concentration was also decreased significantly in trees treated with combination treatment relative to control (*P* < 0.001; Fig 8).

**Fig. 7.**
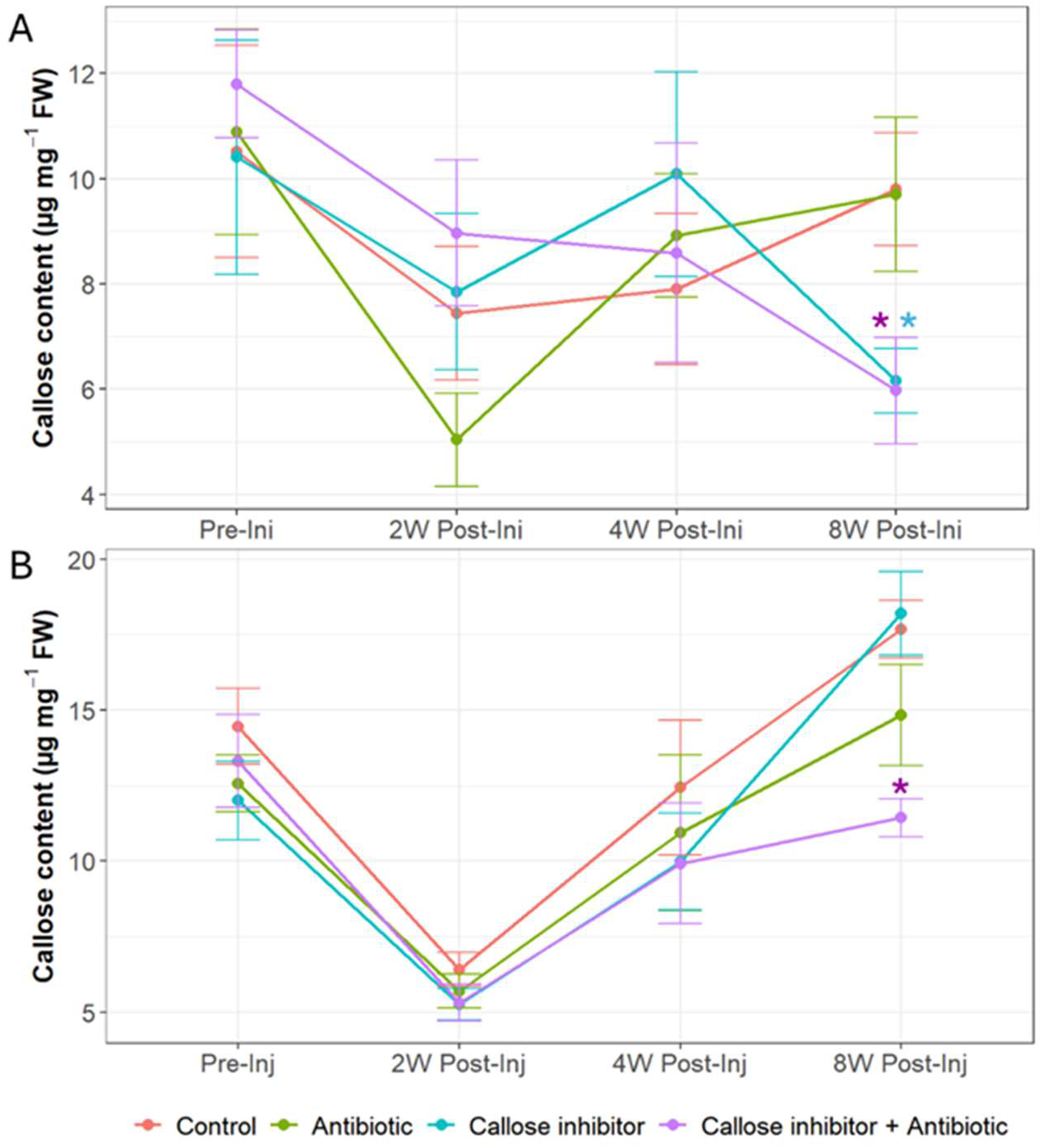
Callose content in leaf midribs of HLB-infected ‘OLL-8’ sweet orange (*C. sinensis*) grafted onto UFR-1 rootstock (*Citrus spp.*), treated with oxytetracycline, the callose inhibitor (3AB), their combination, or left untreated across pre- and three post-injection intervals in (A) 2024 and (B) 2025. Asterisks, colored by treatment, denote significant differences from the control (*P* < 0.05).

**Fig. 8.**
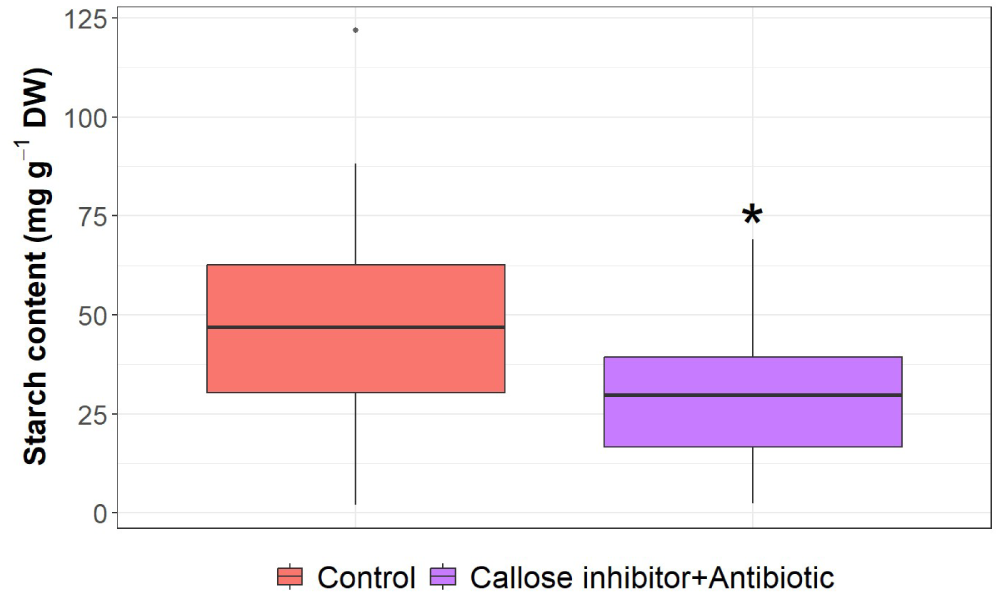
Leaf starch content in HLB-infected ‘OLL-8’ sweet orange (*C. sinensis*) grafted onto UFR-1 rootstock (*Citrus spp.*) under combined oxytetracycline and callose inhibitor (3AB) treatment or left untreated (control). An asterisk denotes a statistically significant difference from the control.

## Discussion

Our results demonstrate that simultaneously targeting the pathogen and the disease-induced callose accumulation can restore phloem function and enhance productivity in citrus trees affected by HLB. These findings support the hypothesis that impaired sugar export and transport in infected trees results from the combined effects of excessive callose deposition and high populations of *C*Las within sink tissues, rather than a single causal factor. While callose-mediated occlusion of sieve elements in infected trees is widely recognized as a primary barrier to phloem transport (Kim et al. 2009), our data indicate that bacterial accumulation at terminal sink pathways imposes an additional significant constraint on assimilate flow. Sink strength is a determinant of phloem mass flow because active unloading and metabolism at sinks maintain the osmotic gradient that drives pressure-flow (Knoblauch et al. 2016; Lemoine et al. 2013). This concept is supported by the mechanistic model of Minchin et al. (1993), which demonstrates that sink strength, defined by unloading kinetics, directly influences phloem pressure gradients and thus governs mass flow and sink priority. Collectively, these studies provide strong independent support for our interpretation that high bacterial populations that impair sink metabolism can act as a terminal bottleneck on sugar flow in HLB, which explains why only the combined antibiotic and callose-inhibitor treatment restored export in our experiments. These findings highlight that effective HLB management requires addressing both pathogen load and host-induced phloem blockage, thereby offering a new integrated approach for mitigating HLB symptoms and sustaining productivity in infected citrus orchards.

Among the treatments evaluated, only the combination of antibiotic and callose inhibitor consistently improved tree health, fruit retention, carbon allocation, and total carbon accumulation in fruits after physiological fruit drop. In contrast, individual applications of either treatment provided only limited or transient benefits. Antibiotic treatment alone can reduce *C*Las titers (Archer et al. 2022; 2023; Hu and Wang 2016; Hu et al. 2018; Roldán et al. 2022; 2024), which may indirectly decrease callose deposition. However, efficacy is inconsistent, influenced by infection stage and cultivar, and yield gains are not always statistically significant (Archer et al. 2023; Hu and Wang 2016). In our study, antibiotic alone temporarily increased carbon import into fruits without improving physiological fruit drop or final fruit carbon accumulation. Although previous studies have reported that OTC trunk injection can reduce preharvest fruit drop and increase yield (Archer et al. 2022; 2023), yield losses in HLB-affected trees are also driven by early physiological fruit drop and failure of fruit set on symptomatic branches (Bassanezi et al. 2011), underscoring the importance of targeting early fruit drop when evaluating treatment effectiveness. During our sampling period, antibiotic treatment alone did not reduce leaf *C*Las titers, suggesting that OTC requires a longer period to reach foliar tissues and achieve detectable pathogen suppression. Consistent with this interpretation, OTC has been shown to accumulate slowly in leaves following trunk injection, reaching peak concentrations approximately 120 days post-injection (Archer et al. 2022), with significant effects on leaf Ct values observed between one and six months (Archer et al. 2023; Vincent et al. 2022). Consequently, although *C*Las suppression may precede physiological recovery, reductions in callose deposition and restoration of phloem function are likely delayed under antibiotic-only treatment. Moreover, systemic antibiotic distribution may depend on phloem integrity, as callose-induced blockage can impede transport by disrupting carbohydrate export and reducing xylem hydraulic conductivity. In contrast, callose synthesis inhibitors facilitate reopening of occluded phloem channels, thereby enhancing OTC translocation and overall treatment effectiveness. Consistent with this, the present study shows that antibiotic treatment alone did not significantly reduce bacterial load, whereas the combined treatment effectively lowered it. Callose inhibition alone provided rapid but short-lived improvements in phloem function, as demonstrated by the ^14^C experiment showing increased sucrose export from source leaves and import into fruits within two weeks of infiltration; however, these benefits were not sustained under long-term field conditions. Lower Ct values observed at two and four weeks after injection suggest that reopening the phloem while bacterial cells were still present may have facilitated pathogen movement within the plant, contributing to increased phloem dysfunction over time and limiting the long-term effectiveness of callose inhibition alone. The pronounced reduction in relative fruit growth for this treatment compared with the control during the four- to six-week post-injection interval further supports this interpretation. Similarly, Li et al. (2016) reported that repeated applications of 2-DDG reduced HLB symptom severity and improved fruit yield and quality, although yield gains were not significantly greater than those of untreated trees. Overall, these results indicate that neither treatment alone is ideal for sustained recovery, highlighting the value of combining pathogen suppression with regulation of host phloem function to achieve lasting phloem recovery and improved productivity in HLB-affected citrus.

Improved carbohydrate transport in the combined OTC-callose inhibitor treatment indicates that phloem function was recovered, probably as a result of decreased callose deposition and lower bacterial load. Under disease-free conditions, callose deposition in sieve pores maintains phloem integrity and seals wounds. However, during biotic stress, excessive and persistent callose can obstruct sieve plates and disrupt source-sink balance (Achor et al. 2020; Koh et al. 2012). Moderation of pathological callose formation through combined chemical inhibition and pathogen suppression can reopen transport pathways and restore assimilate flow. In the first year of the experiment, both the callose inhibitor alone and the combined treatment reduced callose accumulation eight weeks after injection. In the second year, only the combined treatment maintained this reduction at the same time point, indicating more sustained suppression compared with the callose inhibitor alone. Effective phloem recovery therefore depends on continued regulation of both pathogen levels and callose synthesis. Supporting this, the combined treatment also eliminated *C*Las in the leaves at eight weeks post-injection. Lower callose levels coincided with higher Ct values and reduced starch accumulation in the leaf, all reflecting restored phloem conductivity and carbohydrate transport capacity (Etxeberria et al. 2009). Although phloem function appeared to recover, visible canopy growth remained largely unchanged, suggesting that external recovery lags behind internal physiological restoration. However, trees receiving the combined callose inhibitor and antibiotic treatment showed a moderate increase in canopy density, indicating improved carbon distribution and renewed leaf growth. While this increase was not statistically significant at the conventional threshold (*P* = 0.06), the trend suggests a gradual re-establishment of source-sink coordination. Notably, following hurricane-induced defoliation, trees treated with the callose inhibitor, or the combination treatment exhibited greater canopy resilience. Thus, canopy density, rather than canopy volume, may more accurately reflect physiological recovery, consistent with previous observations that HLB-affected trees can regain productivity without substantial changes in overall size (Levy et al. 2023).

Despite minimal changes in overall tree growth, improved phloem conductance enhanced the delivery of photoassimilates to developing fruits during periods of high sink demand. Trees receiving the combined antibiotic and callose inhibitor exhibited increased fruit growth rates at two weeks and again at six to eight weeks post-treatment. Correspondingly, estimated carbon import into the developing fruit at four- and eight-weeks post-injection was higher in this treatment group. These intervals coincide with key stages of fruit development, particularly the physiological fruit drop period (May–June drop; Banjare et al. 2023), when assimilate competition is high (Dovis et al. 2014; Iglesias et al. 2003; Iglesias et al. 2007; Li et al. 2017). During this stage, carbohydrate shortage in developing fruitlets triggers hormonal imbalances, elevated abscisic acid and ethylene and reduced auxin and gibberellin activity, that promote abscission (Gómez-Cadenas et al. 2000; Iglesias et al. 2006; Kuang et al. 2012). Interestingly, the combined treatment reduced physiological fruit drop, likely by stabilizing carbohydrate availability and hormonal signaling through enhanced assimilate transport. This treatment also exhibited elevated fruit respiration at both sampling times, suggesting that additional assimilates supported energy-demanding stress responses against HLB, including phloem recovery and defense mechanisms that promote fruit retention despite the higher carbon cost per fruit. Consequently, the combined treatment may improve yield by reducing fruit abscission. Consistent with this, the callose inhibitor plus antibiotic treatment increased the total carbon accumulated in the fruits after physiological fruit drop relative to the control.

Taken together, these findings demonstrate that simultaneously targeting the pathogen and callose-mediated phloem blockage is more effective than addressing either factor alone, which we explain by proposing a source-conduit-sink model of carbon transport dysfunction and recovery in HLB-affected citrus (Illustrated in Fig 9). Under healthy conditions, photoassimilates produced in source leaves are loaded into the phloem and transported through low-resistance sieve elements toward strong sinks, such as developing fruits, where active unloading and metabolism maintain sink-driven pressure gradients. When affected by HLB, pathogen-induced callose accumulation narrows or blocks sieve pores, increasing resistance within the phloem conduit, while bacterial proliferation in sink tissues impairs unloading capacity and weakens sink strength. Together, these effects restrict transport along the pathway and create a terminal bottleneck at the sink, reducing effective assimilate flow despite continued source activity.

**Fig. 9.**
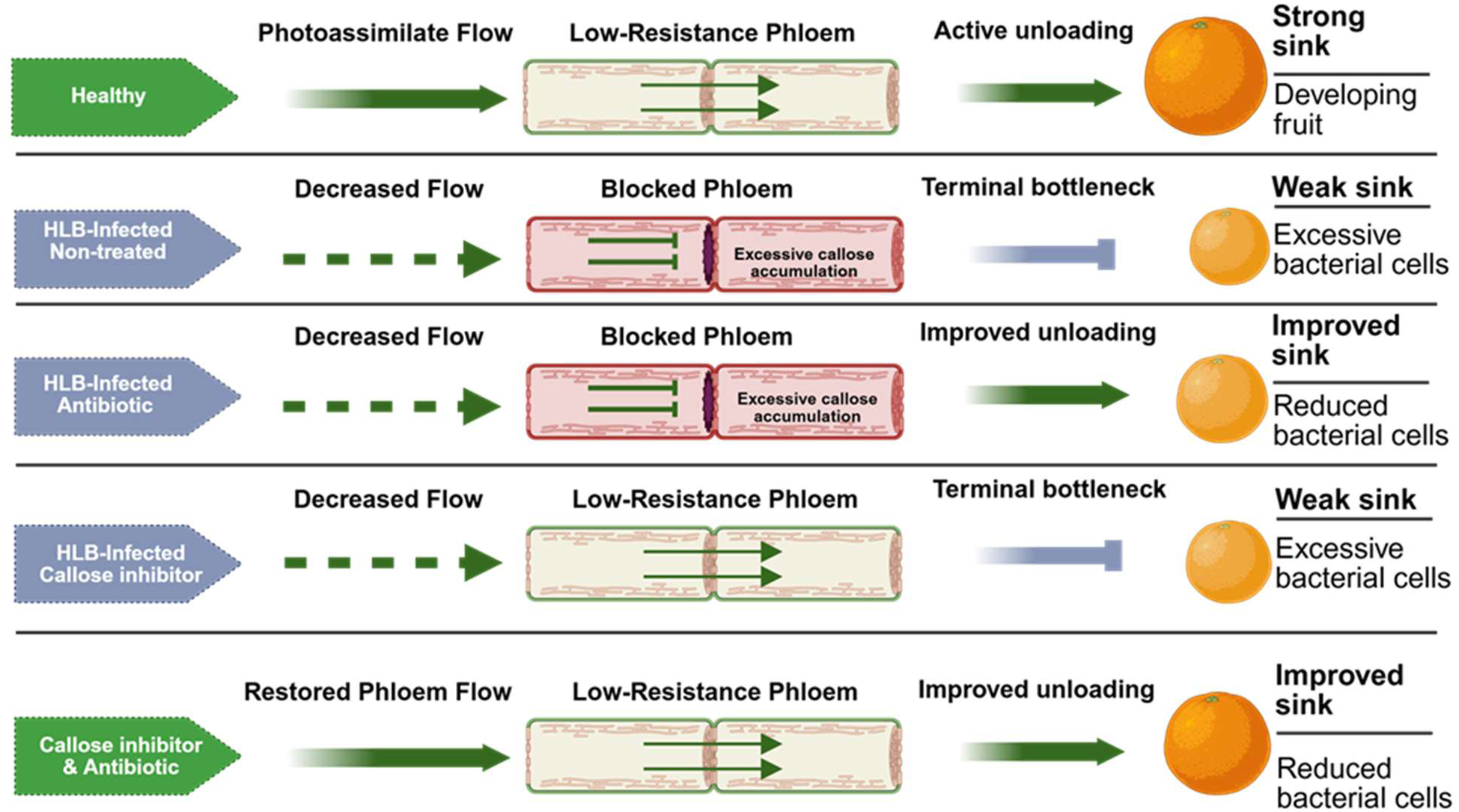
Conceptual source-conduit-sink model of carbon transport dysfunction and recovery in HLB-affected citrus. Image created with BioRender (www.biorender.com)

Application of a callose inhibitor alone reopens sieve elements and transiently improves conductivity but does not restore sustained transport in the presence of dysfunctional sinks. Conversely, antibiotic treatment alone reduces bacterial load and partially improves sink function but remains limited by persistent conduit obstruction. Only the combined treatment restores both conduit integrity and sink activity, allowing pressure-driven mass flow to resume, enhancing carbon import into fruits, reducing fruit abscission, and improving yield (Fig 9). This model integrates our physiological and growth results and provides a mechanistic basis for the observed synergistic treatment effects. By improving phloem transport and maintaining carbon supply during periods of high sink demand, this combined approach helps support fruit retention and overall productivity in HLB-affected citrus. Future work should focus on optimizing treatment dosage, identifying optimal seasonal timing, and assessing how different citrus cultivars respond to this strategy under field conditions.

## Acknowledgements

Thank you to Myrtho Pierre, Gillian Zeng Michalczyk, Dorcas Alade, Rashmi Kumari, Muna Aryal, Jean Francois and Donielle Turner for all the help and support during the data collection process, Nabil Killiny for his input on antibiotic use and experimental design, and Jason Griffin for the use of his trees for the short-term experiment. This work was supported by the Citrus Research and Development Foundation (CRDF) project number 22-017.

## SUPPLEMENTARY MATERIAL

**Supplementary Figure 1.**
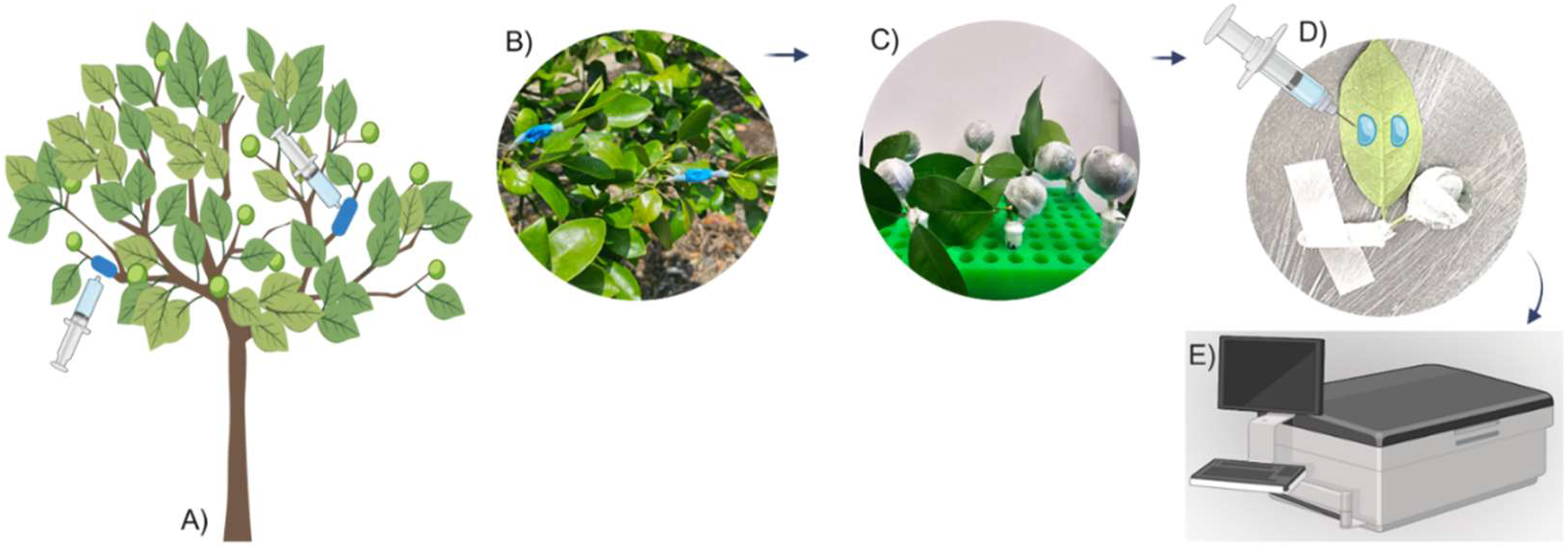
Overview of the methodology for short-term experiment: from tree treatment infiltration (A and B) to sucrose ^14^C labeling (C and D) and Liquid Scintillation detection (E).

**Supplementary Figure 2.**
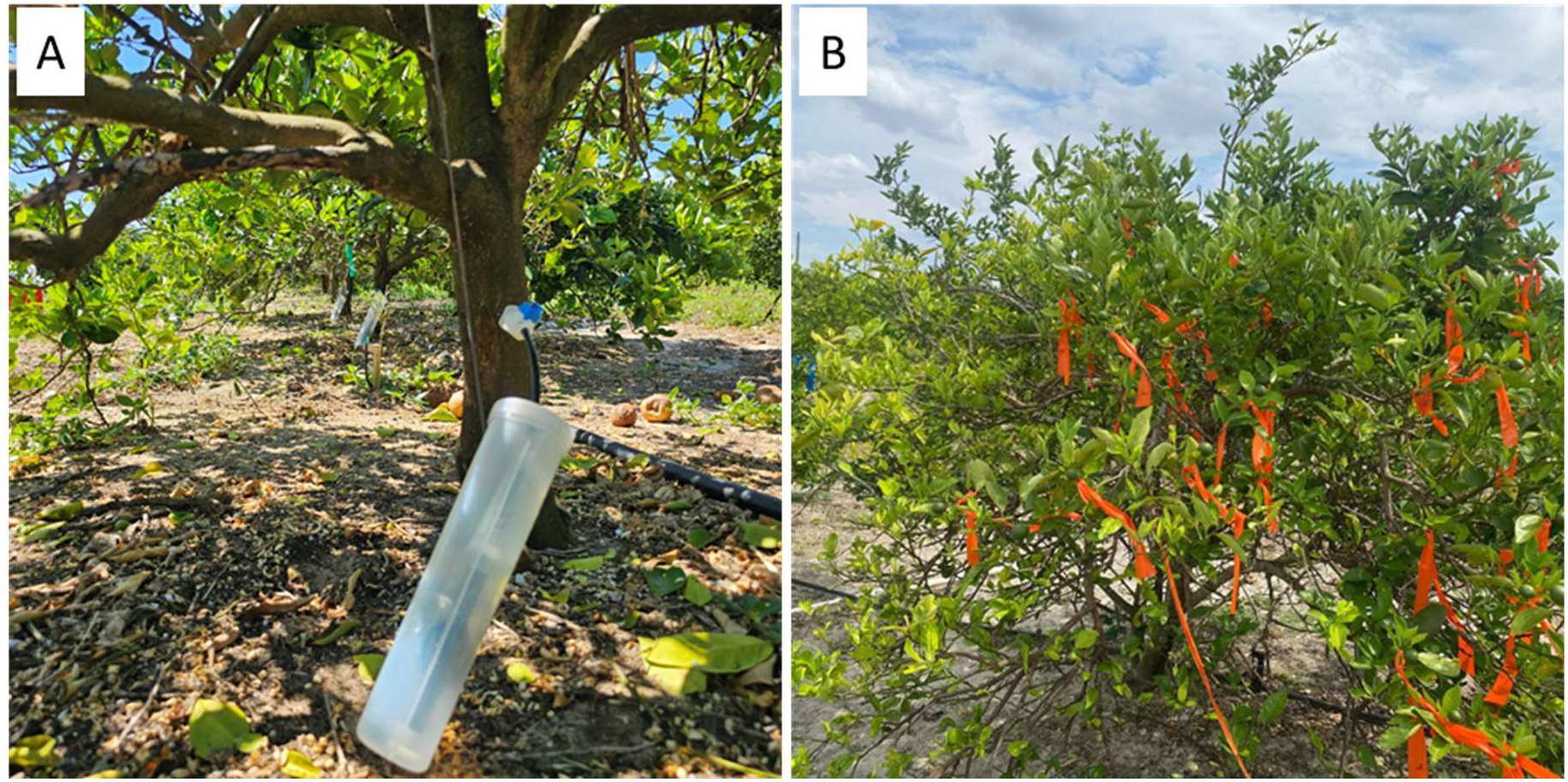
Trunk injection procedure and fruit tagging. (A) Trunk injection of the prepared solution, with each tree receiving 50 mL, delivered on the north side of the trunk using a large FlexInject™ injector fitted with a 17/64-inch sharp brad-point drill bit. (B) Representative view of tagged fruits; on each tree, 30 fruits were selected and labeled for subsequent evaluation.

## Notes

### Competing Interest Statement

The authors have declared no competing interest.

